# A time-dependent mechano-bioenergetics model of muscle contraction

**DOI:** 10.64898/2026.06.24.734405

**Authors:** R. N. Konno, G. A. Lichtwark, T. J. M. Dick

## Abstract

Predictions of skeletal muscle energy consumption under a diverse range of muscle contractile conditions are critical for improving our understanding of locomotion. Existing mathematical models, while capturing the mechanical dependence of energy consuming processes, neglect the time-dependent behaviour and recovery costs associated with regenerating ATP. This time-dependence is important for predicting the energetic response of muscles during repetitive or cyclical tasks like locomotion, where muscle undergoes many contraction cycles. This study presents a novel model to predict energetic rates based on physiological processes: Ca^2+^ transport costs, cross-bridge cycling costs, and ATP regeneration. Previous mathematical models include the dependence on Ca^2+^ transport and cross-bridge cycling, but neglect the time-dependent response and the subsequent recovery of ATP following the contraction. Model parameters were obtained from existing data on isolated muscle preparations, and predicted energetic rates were validated on separate datasets across a range of contractile conditions including dynamic, sub-maximal, and twitch contractions. The time-dependent model was able to capture the influence of contraction frequency on peak energetic rates and the time-course of energetic recovery observed experimentally. The model captures key physiological processes while maintaining a minimal number of free parameters and low computational cost. This enables generalisability across muscles and species, and implementation into larger scale musculoskeletal models.

## 1 Introduction

Estimates of skeletal muscle energy consumption over the course of a contraction cycle are critical for understanding a muscle’s function during any locomotor task and for predicting the energetic costs of movement. While it is possible to experimentally obtain direct measures of energy use *in vivo* (averaged over many seconds (Haeufle et al., 2020) or minutes of steady-state conditions (Cavagna et al., 1977)), isolated *ex vivo* muscle preparations are required to obtain the precise time-dependent energetic responses of muscle (Woledge et al., 1985). In many movements, such as walking or running, muscles undergo repeated cyclical contractions, meaning that averaged values are not capable of giving us insight into the relationship between muscle mechanical state (muscle strain, strain rate, and activation level) and energy use; thus, mathematical models are required to estimate these relationships. However, existing models (e.g., Konno et al. (2025) and Lichtwark and Wilson (2005)) rely predominantly on the mechanical state, and neglect the muscle history and recovery processes that are also key determinants of muscle energy use (Barclay & Curtin, 2023; Lichtwark et al., 2025).

Energy, in the form of ATP, is consumed during a muscle contraction to transport Ca^2+^ into the sarcoplasmic reticulum and to cycle the actin-myosin cross-bridges. Both of these processes are dependent on the mechanical state of the muscle (Woledge et al., 1985), and as muscle shortens at faster rates it produces more heat (Fenn, 1923). Existing energy models account for these contributors (Umberger et al., 2003), but critically, are not modelled based on the true energy consuming processes. Instead, the activation heat associated with Ca^2+^ transport is typically calculated as proportional to the level of activation and not to the rate of Ca^2+^ transport (Konno et al., 2025). Further, given that the rate of Ca^2+^ transport depends on the concentration of Ca^2+^ within the cytoplasm of muscle fibres, current models cannot account for the history dependence on the rate of Ca^2+^ transport.

As ATP is the fuel consumed during the muscle contraction, its supply needs to be regenerated through either aerobic or anaerobic processes. These regeneration mechanisms are not completely efficient, and heat is released during the conversion of the energy stored in substrates (i.e., glucose or palmitate) to ATP (Barclay, 2019). These energy costs are of similar magnitude to the initial costs of generating force (Barclay, 2019), and are thus likely an important contributor to total *in vivo* energy use, particularly during repeated contractions or longer durations. While existing models captured the time-course of ATP regeneration (Kushmerick, 1998), they have not been coupled with models that capture the initial consumption of ATP, nor the inefficiencies in this process, limiting their ability to capture whole muscle energy use.

In this study, we present and validate a novel mechano-bioenergetics model accounting for the time-dependent processes governing the energetics of muscle contraction, including both the initial mechanisms capturing ATP consumption and the subsequent regeneration of ATP. The initial contractile energetics model is based on Konno et al. (2025), accounting for the mechanical dependence of muscle energetic rates, with modifications to directly model Ca^2+^ transport costs. To capture the time-course of ATP recovery, we implemented a bioenergetics model based on Kushmerick (1998) with an additional component accounting for the inefficiencies of the regeneration process. The model parameters are informed using data from isolated mouse muscle preparations (Barclay et al., 2010; Barclay et al., 1995), and validated on separate datasets across a range of contractile conditions and stimulation intensities (Barclay, 2012; Barclay & Weber, 2004; Lewis & Barclay, 2014; Mast & Elzinga, 1987, 1988).

## 2 Methods

### 2.1 Mathematical model

The modelling framework consists of a combination of individual models capturing the process of muscle contraction (Fig. 1): i) the excitation-activation model captures the flow of Ca^2+^ upon stimulus and the subsequent binding of Ca^2+^ to troponin, ii) the mechanical Hill-type model simulates the muscle mechanical state (strain, strain rate, and force), iii) the initial energetics model captures the energetic cost associated with Ca^2+^ pumping into the sarcoplasmic reticulum and cross-bridge cycling, and iv) the bioenergetics model captures the recovery processes including the ATP-PCr dynamics and the corresponding energetic rate.

**Figure 1:**
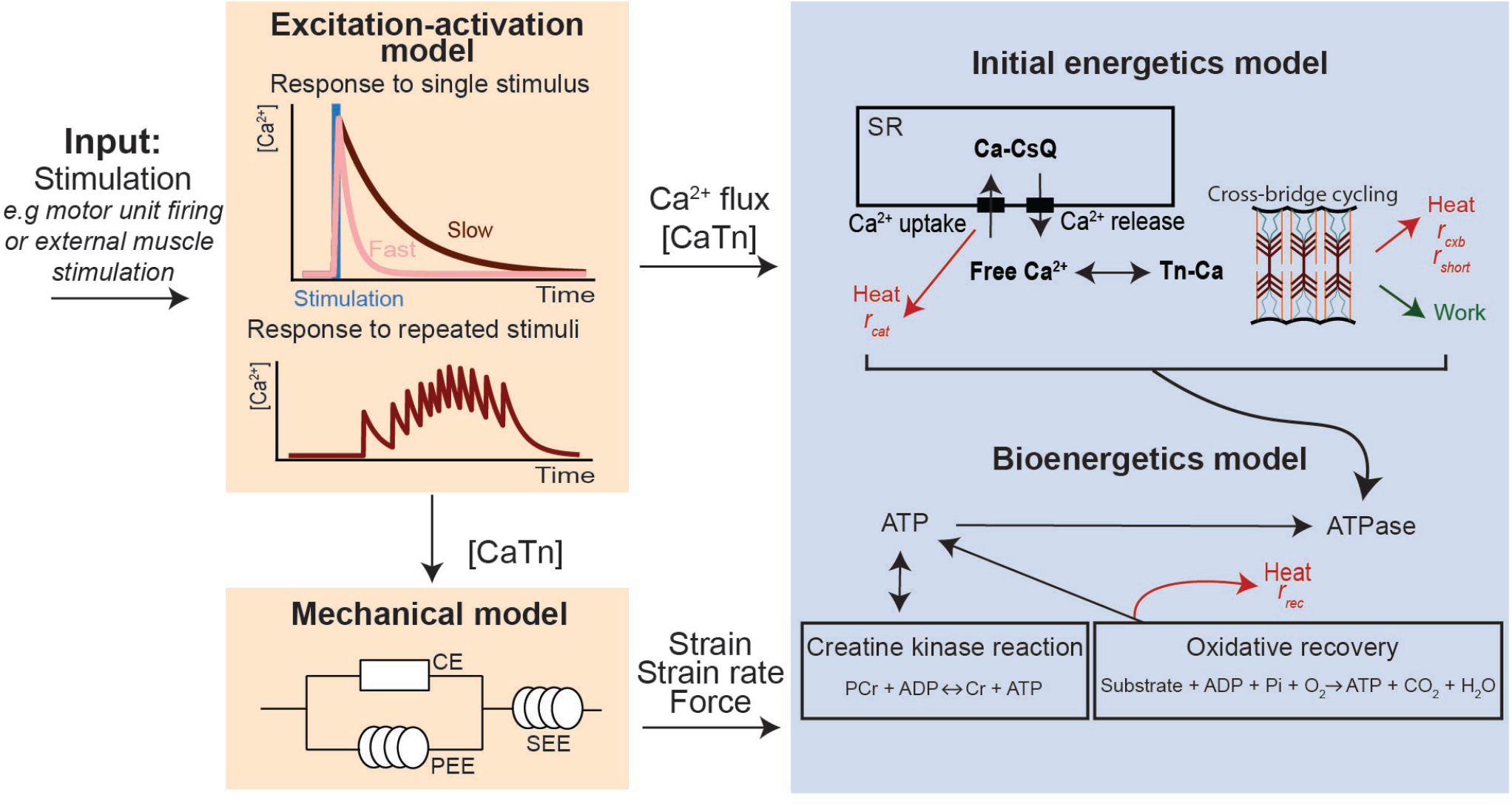
Model workflow. The mathematical model takes as input a stimulus, either a motor unit firing or external stimulation. From the stimulus, the excitation-activation model computes the Ca^2+^ concentrations and the activation state (∝ [CaTn]). The activation state is used by the mechanical model to solve for the muscle strain, strain rate, and force. The outputs from these two models are then used by the initial energetics model to determine the energy used during the contraction, which gives the ATP consumed by ATPases. The initial energetics model computes the heat lost as inefficiencies to Ca^2+^ transport (parameter r_cat_), cross-bridge cycling (parameter r_cxb_), and additional heat lost to inefficiencies during shortening (parameter r_short_). The regeneration of ATP consumed is determined using a bioenergetics ATP-PCr model, which solves for the ATP and PCr levels within the muscle. The rate of regeneration of ATP is calculated as a function of the ADP within the muscle. Here we assume oxidative recovery, and the heat released during regeneration of ATP is proportional (parameter r_rec_) to the rate of regeneration of ATP. CE: contractile element, PEE: parallel elastic element, SEE: series elastic element, SR: sarcoplasmic reticulum, CsQ: calsequestrin, Tn: troponin, PCr: phosphocreatine, Cr: creatine, ATP: adenosine triphosphate, ADP adenosine diphosphate, Pi: inorganic phosphate.

#### 2.1.1 Excitation-activation model

Upon a stimulus, Ca^2+^ is released from the sarcoplasmic reticulum into the cytoplasm. Here, we use a simplified model to limit the unknown parameters required to capture *in vivo* muscle contraction (Fig. 1). We account for the troponin bound Ca^2+^, which is proportional to cross-bridge cycling and force production; thus, the Ca^2+^ -troponin complex gives a measure of muscle activation. After stimulus, ATP is required to pump the Ca^2+^ against the concentration gradient back into the sarcoplasmic reticulum. We capture the flow of Ca^2+^ ions throughout the membrane based on an existing model (Mayfield et al., 2023)

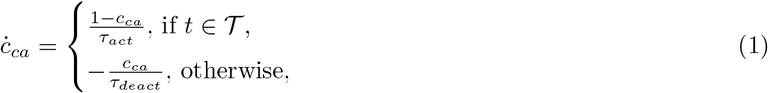

where 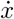 denotes the time derivative of a given quantity *x, t* is the time, *c*_*ca*_ is the free Ca^2+^ concentration within the cytoplasm, and *τ*_*act*_ and *τ*_*deact*_ are the activation (release of Ca^2+^) and deactivation (uptake of Ca^2+^) time constants, respectively. *T* denotes the time period where the muscle is stimulated. The concentration of bound Ca^2+^ to troponin is calculated as

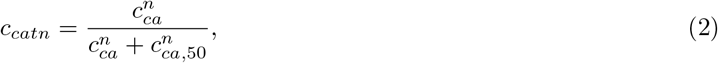

where *n* is the Hill constant and *c*_*ca*,50_ is the concentration of calcium at half of the maximal cross-bridge activation.

#### 2.1.2 Mechanical model

As we follow the formulation of the muscle mechanical model presented elsewhere (Dick et al., 2017; Konno et al., 2025), the details are contained in the Supplementary Material. The Hill-type model was used to compute estimates of muscle strain, strain rate, and force based on the activation level predicted by the excitation-activation model (Fig. 1). We assume that muscle activation is proportional to the Ca^2+^ bound to troponin, and since we compute a normalised 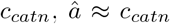 Normalised muscle force was calculated based on a Hill-type model

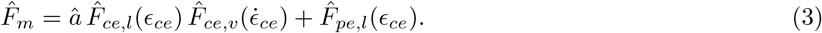

Here, 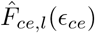 is the strain dependence of active force production, 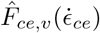 is the strain rate dependence of the active force production, and 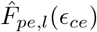 is the passive response. *ϵ*_*ce*_ is the strain in the contractile element computed as *ϵ*_*ce*_ = (*l − l*_0_)*/l*_0_, where *l* is the current length of the contractile element and *l*_0_ is optimal length.

#### 2.1.3 Initial energetics model

Energetic cost from the initial phase of the contraction (Ca^2+^ transport and cross-bridge cycling), was modelled following existing models (Konno et al., 2025; Lichtwark & Wilson, 2005), but adapted to capture the energetic cost associated directly with Ca^2+^ transport and binding based on the Ca^2+^ model. We start by defining the isometric cross-bridge heat rate as

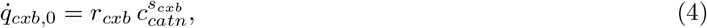

where *r*_*cxb*_ gives the energetic constant of cross-bridge cycling and *s*_*cxb*_ is a scale factor to account for the Ca^2+^ dependence of the cross-bridge energetic rates (Lewis & Barclay, 2014). To account for the length dependence of the maintenance heat rate, we scale the the isometric rate by the force-length relationship (Homsher et al., 1972) giving us

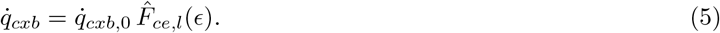

The energy consumed due to Ca^2+^ transport is proportional to the rate of Ca^2+^ pumping into the sarcoplasmic reticulum. Note, we assume negligible Ca^2+^ transport heat during the stimulation period as the duration of the stimulus is small. The Ca^2+^ transport heat rate is given as

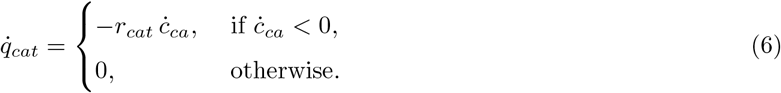

*r*_*cat*_ is the energetic constant associated with Ca^2+^ transport.

We account for the dependence of muscle heat rate on the muscle mechanical state through the shortening and lengthening heat rates, which are characterised in Konno et al. (2025). We define the mechanical heat rate dependence as

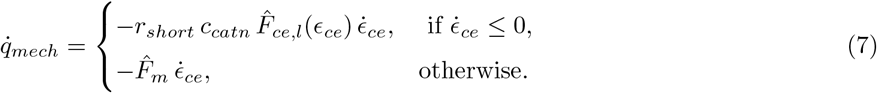

*r*_*short*_ is the energetic constant of shortening. Note that if lengthening 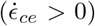, we assume any work done on the muscle is eventually released as heat (Linari et al., 2003). The work rate performed by the muscle is defined as

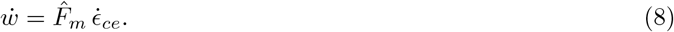

The total initial energetic rate is then assumed to be a combination of the above rates

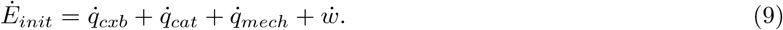

The total initial energy predicted here must be recovered through the ATP recovery processes, thus, these energy rates are used as input to the bioenergetics model.

#### 2.1.4 Bioenergetics model

The amount of ATP consumed through the initial phase of muscle contraction needs to be regenerated through metabolic pathways. Here we assume the relationship between ATP flux, *ϕ*_*ATP*_, and the initial energy consumed to be given by

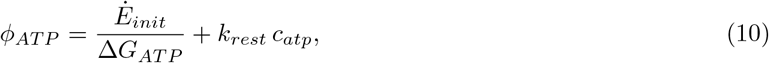

where Δ*G*_*ATP*_ = 60 *×*10^3^*J mol*^*−*1^ is the free energy available during hydrolysis of ATP (Barclay, 2019). *k*_*rest*_ *c*_*atp*_ accounts for resting state ATP consumption.

This bioenergetics model presented here is based on an existing formulation (Kushmerick, 1998; Vicini & Kushmerick, 2000). The concentration of ATP, *c*_*atp*_, and PCr, *c*_*pcr*_, can be written in terms of the flux of ATP consumed, *ϕ*_*ATP*_, and ATP regeneration by aerobic pathways (*ϕ*_*ox*_) and creatine kinase (*ϕ*_*CK*_). The system of differential equations along with initial conditions is

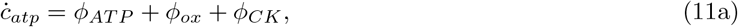

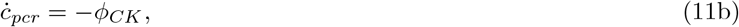

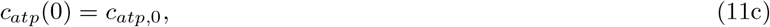

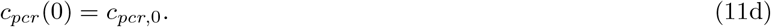

To ensure a balance of adenosine and creatine, we enforce

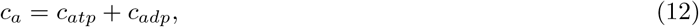

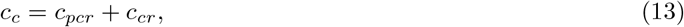

where *c*_*a*_ is the total concentration of adenosine, *c*_*adp*_ is the concentration of ADP, *c*_*c*_ is the total concentration of creatine, and *c*_*cr*_ is the concentration of free creatine.

We assume the regeneration of ATP is largely due to oxidative phosphorylation and neglect the role of glycolysis in influencing muscle pH and the associated consequences (Lambeth & Kushmerick, 2002). While numerous models have been developed to capture the oxidative phosphorylation response as a function of either PCr or ADP (Kushmerick, 1998), we follow Vicini and Kushmerick (2000) and utilise a dependence on ADP as this sufficiently captures work-relaxation relationships (Kushmerick, 1998). Thus, *ϕ*_*ox*_ is given by

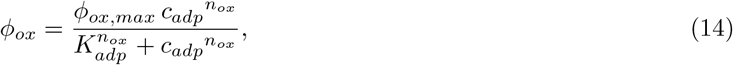

where *ϕ*_*ox,max*_ is the maximum rate of oxidative phosphorylation, *K*_*adp*_ is the concentration of ADP at a rate of half of *ϕ*_*ox,max*_, and *n*_*ox*_ is the Hill coefficient.

The creatine kinase term accounts for the flux between PCr and Cr, and is modelled as

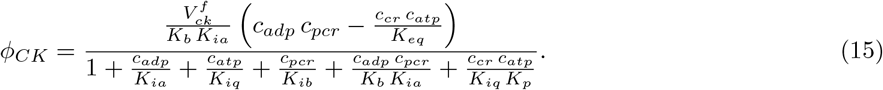

The constants *k*_*b*_, *K*_*ai*_, *K*_*bi*_, *K*_*iq*_, *K*_*p*_ are the rate constants (Vicini & Kushmerick, 2000), while *K*_*eq*_ is the equi-librium constant (Lawson & Veech, 1979). 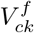is the forward rate creatine kinase. This relationship is obtained from Vicini and Kushmerick (2000), equation 5, assuming that reverse rate of creatine kinase is proportional to the forward rate

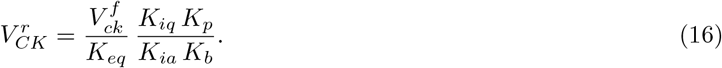

To determine initial concentrations of ADP and the resting rate coefficient, we use the steady-state equations by setting *ϕ*_*CK*_ = 0 and *ϕ*_*atp*_ = *ϕ*_*ox*_. This gives

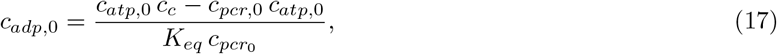

and

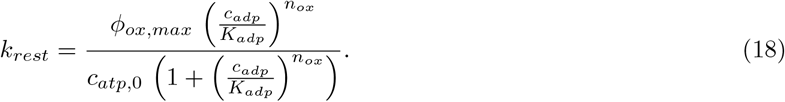

By solving equations Eqs.(11a)-(11d) we obtain the rate of oxidative phosphorylation, which, if the efficiency of the recovery process is known, we can compute the energetic rate of the recovery processes

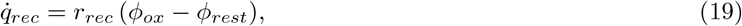

where *r*_*rec*_ is the recovery heat constant. As we are interested in only the energy consumed as a result of the contraction process, we subtract of the ATP flux due to resting processes, *ϕ*_*rest*_ = *k*_*rest*_ *c*_*atp*,0_. A summary of the all parameters (values constant within the model) and variables (values changing with time) used in the models are provided in Table 1 and Table 2.

**Table 1:**
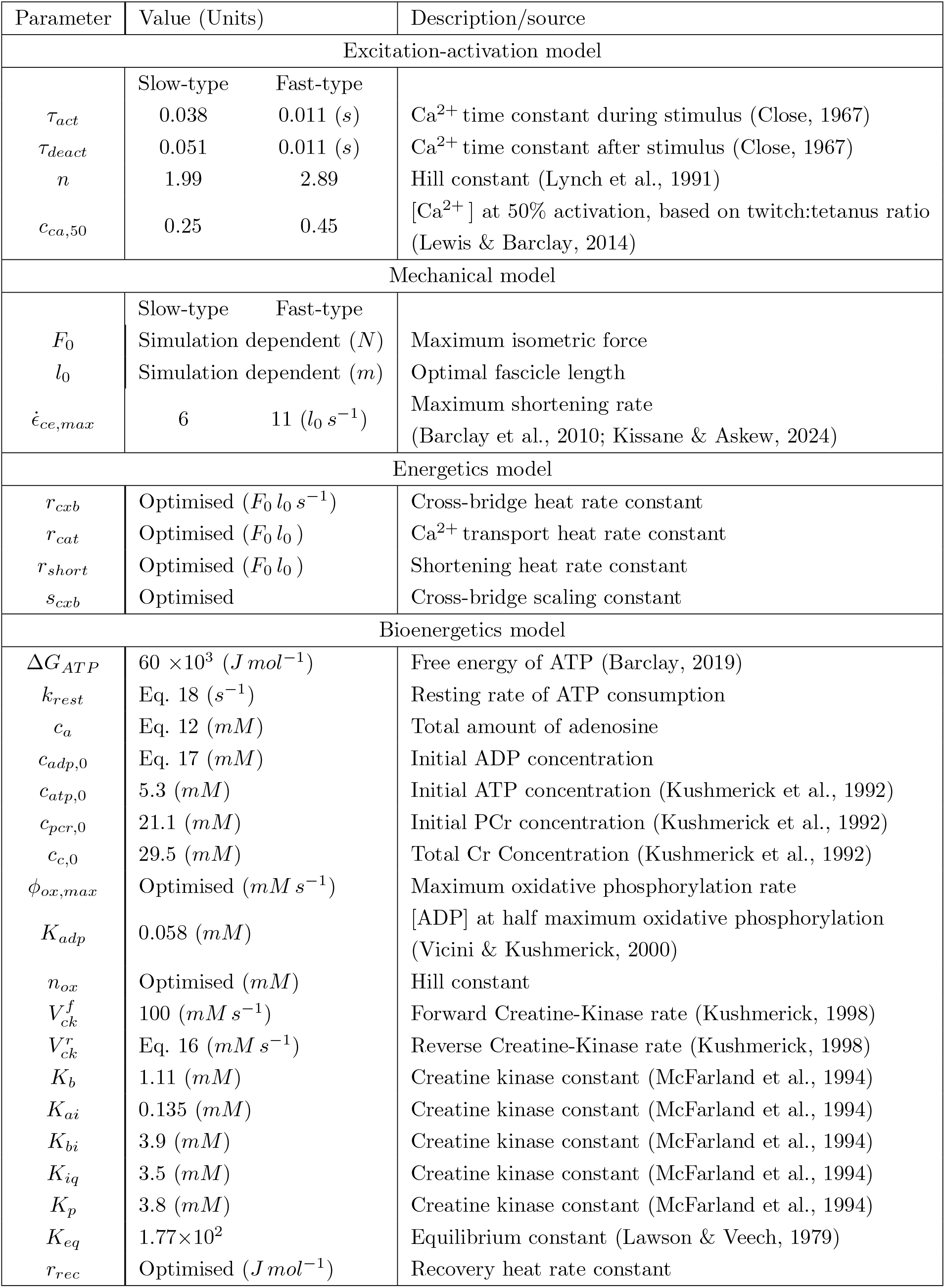
Parameters included in the mechano-bioenergetics model, along with their source if available. If no units are given then the parameter is either normalised with normalisation values given or a unitless parameter.

**Table 2:**
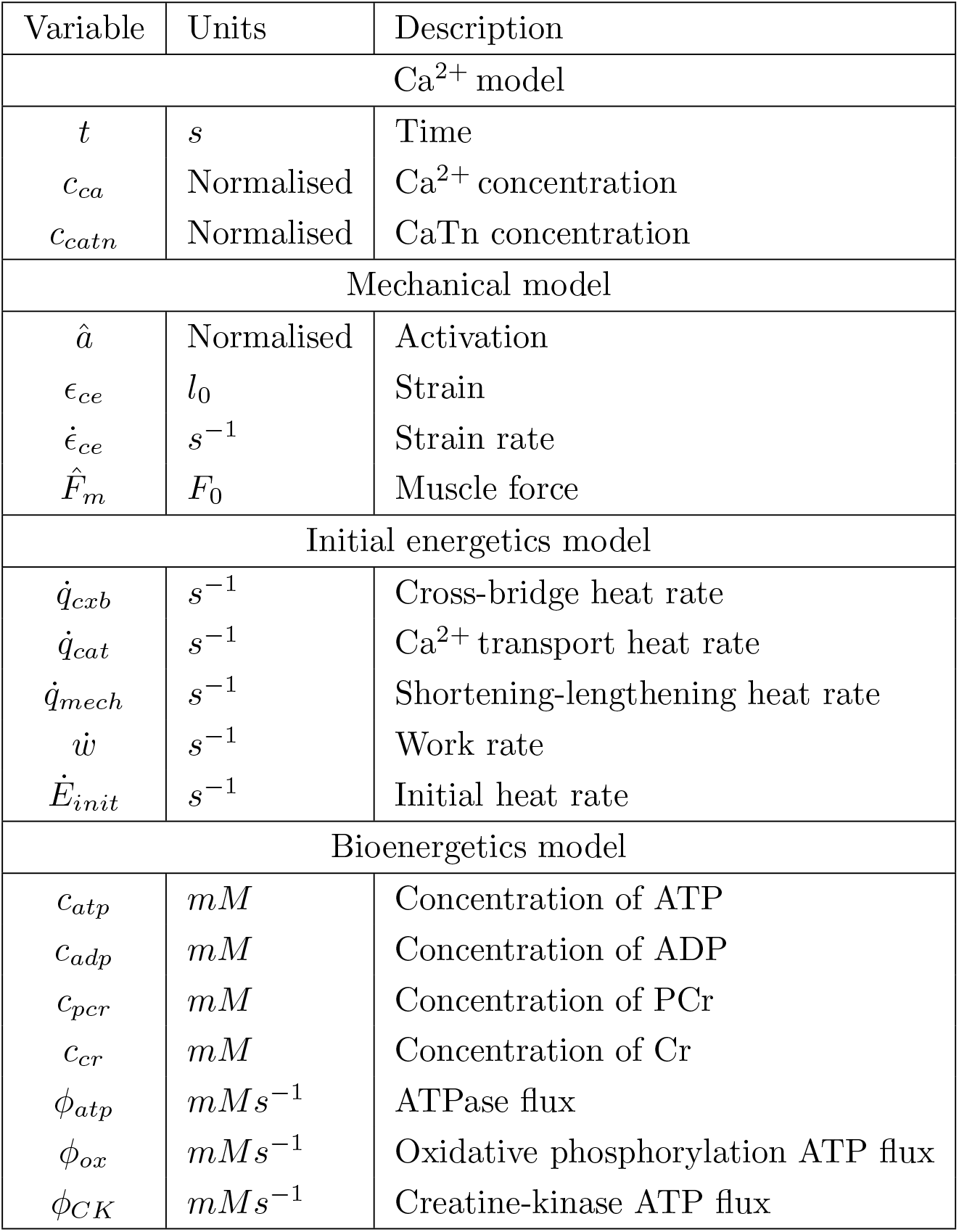
Variables and functions included in the bioenergetics model along with units and description.

| Variable | Units | Description |
| --- | --- | --- |
| Ca<sup>2+</sup> model |  |  |
| $t$ | $s$ | Time |
| $c_{ca}$ | Normalised | Ca<sup>2+</sup> concentration |
| $c_{catn}$ | Normalised | CaTn concentration |
| Mechanical model |  |  |
| $\hat{a}$ | Normalised | Activation |
| $\epsilon_{ce}$ | $l_0$ | Strain |
| $\dot{\epsilon}_{ce}$ | $s^{-1}$ | Strain rate |
| $\hat{F}_m$ | $F_0$ | Muscle force |
| Initial energetics model |  |  |
| $\dot{q}_{cxb}$ | $s^{-1}$ | Cross-bridge heat rate |
| $\dot{q}_{cat}$ | $s^{-1}$ | Ca<sup>2+</sup> transport heat rate |
| $\dot{q}_{mech}$ | $s^{-1}$ | Shortening-lengthening heat rate |
| $\dot{w}$ | $s^{-1}$ | Work rate |
| $\dot{E}_{init}$ | $s^{-1}$ | Initial heat rate |
| Bioenergetics model |  |  |
| $c_{atp}$ | $mM$ | Concentration of ATP |
| $c_{adp}$ | $mM$ | Concentration of ADP |
| $c_{pcr}$ | $mM$ | Concentration of PCr |
| $c_{cr}$ | $mM$ | Concentration of Cr |
| $\phi_{atp}$ | $mMs^{-1}$ | ATPase flux |
| $\phi_{ox}$ | $mMs^{-1}$ | Oxidative phosphorylation ATP flux |
| $\phi_{CK}$ | $mMs^{-1}$ | Creatine-kinase ATP flux |

### 2.2 Model Parameters

The excitation-activation model parameters were chosen based on experimental data from isolated muscle experiments (see Table 1). The parameters of the Hill-type model have been described previously (Dick et al., 2017) with adjustments made for 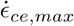 for mouse muscle fibres (Barclay et al., 2010; Kissane & Askew, 2024). The initial energetics parameters (*r*_*cxb*_, *r*_*cat*_, *r*_*short*_, and *s*_*cxb*_) were determined through an optimisation to the dataset from Barclay et al. (2010). The recovery energetic rate, *r*_*rec*_, and the bioenergetic recovery model parameters, *n*_*ox*_ and *ϕ*_*ox,max*_, were optimised to match the experimental time-course of recovery (Barclay et al., 1995) and subsequently corrected to match the magnitude of the recovery rates with changes to temperature (Barclay, 2019; Barclay & Weber, 2004). A detailed description of the experimental protocols are within the Supplementary Material.

#### Initial energetics parameters

The initial energetics model parameters were obtained based on data in the mouse soleus and extensor digitorum longus (EDL) from Barclay et al. (2010), consisting of energetic rates measured at shortening velocities from isometric to maximum strain rates (5.2*s*^*−*1^ for soleus and 8.9*s*^*−*1^ for the EDL). We assumed that the relative contributions from isometric heat production are split such that 60% is due to cross-bridge cycling and 40% is due to Ca^2+^ transport (Ca^2+^ transport contributions typically vary between 25 to 45% (Barclay & Curtin, 2023)). The increase in energetic cost with faster shortening rates (Fenn, 1923) is given by the shortening heat component. To obtain *s*_*cxb*_, we enforced that the ratio between cross-bridge cycling and Ca^2+^ transport heat was maintained across stimulation frequencies (Lewis & Barclay, 2014). Thus, we minimised the objective function

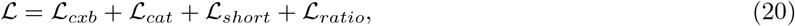

where *L*_*cxb*_ and *L*_*cat*_ enforce the cross-bridge and Ca^2+^ transport heat components, respectively. *L*_*short*_ enforces the mechanical dependence on the shortening rate. *L*_*ratio*_ enforces the ratio in Ca^2+^ transport heat to total initial heat across stimulation frequencies. Objective function details are contained within the Supplementary Material.

#### Bioenergetics model parameters

The bioenergetics model parameters *{r*_*rec*_, *n*_*ox*_, *ϕ*_*ox,max*_*}* were optimised to the experimental data set from (Barclay et al., 1995), where muscle underwent repeated cyclical contractions with a contraction and rest phase. The experimentally determined initial heat rates were used as input to the bioenergetics model, which makes this model independent of the parameters determined in the initial energetics model. The experimental force traces for the EDL demonstrated fatigue, potentially leading to greater inefficiencies in the recovery process not representative of aerobic recovery; thus, we only optimised the parameters to the soleus dataset and applied these parameters to the EDL. The parameters were obtained through minimising the difference between the experimental recovery heat per cycle, 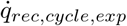, and the recovery heat per cycle predicted by the model, 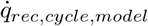,

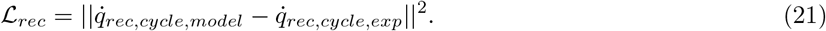

### 2.3 Validation

To validate the model predictions under independent contractile conditions to those in which we developed our model (Barclay et al., 2010; Barclay & Weber, 2004), we compared model outputs to experimental data from across a range of contractile conditions and stimulation frequencies (twitch to tetanus) from Barclay and Weber (2004), Barclay (2012), Lewis and Barclay (2014), Mast and Elzinga (1987), and Mast and Elzinga (1988). To assess both the initial and bioenergetic models, we compared to experimental data for repeated dynamic contractions where the time-course of recovery was measured (Barclay & Weber, 2004). These conditions consisted of repeated contractions at contraction frequencies from 0.5 to 4*Hz* with an overlapping stimulation and shortening phase with the shortening rate chosen to optimise muscle work. Another validation was performed to ensure the time-course captured the energetics when measured using oxygen measures (rather than heat measures). This was done by comparing the time-course of recovery with the experimentally measured time-course following consecutive twitches (Mast & Elzinga, 1987). To assess model performance during sub-maximal contractions (more representative of *in vivo* muscle behaviour), we compared the energy expenditure across a range of stimulation frequencies (Lewis & Barclay, 2014) and consecutive twitches (Mast & Elzinga, 1988), as well as doublet twitches at varying frequencies (Barclay, 2012). A detailed description of the experimental protocols are within the Supplementary Material and all codes are available at https://github.com/ryankonno/TimeDependentEnergetics.

## 3 Results

### 3.1 Parameter estimates

The optimised parameters for the initial energetics and bioenergetics models are given in Table 3 and Table 4, respectively. While the data used to inform the initial energetics model parameters were determined at 30^*°*^C, the bioenergetics data was measured at 20^*°*^C. Thus, as the maximum rate of oxidative phosphorylation scales with temperature (Davies & Tribe, 1969), after the optimisation of the recovery rates (Fig. 2), *ϕ*_*ox,max*_ was scaled by a factor of 2 to give similar time constants to those observed in Barclay and Weber (2004) (Table 4). The ratio of recovery to initial heat was double the values observed in numerous other studies (Barclay, 2019), so we scaled *r*_*rec*_ by half to a value of 0.0825 *J µmol*^*−*1^.

**Table 3:** Optimal parameters for the initial energetics model of the muscle. r_cxb_, r_cat_, r_short_ are the cross-bridge, Ca^2+^ transport, and shortening heat rate coefficients, respectively. s_cxb_ is an exponent used to account for the Ca^2+^ dependence of the cross-bridge heat rate. EDL: extensor digitorum longus.

| Fibre-type | $r_{cxb} (W(F_0 l_0)^{-1})$ | $r_{cat} (J(F_0 l_0)^{-1})$ | $r_{short} (J(F_0 l_0)^{-1})$ | $s_{cxb}$ |
| --- | --- | --- | --- | --- |
| Slow-type (Mouse soleus) | 0.247 | 0.0295 | 0.126 | 0.567 |
| Fast-type (Mouse EDL) | 0.763 | 0.0199 | 0.105 | 0.257 |

**Table 4:** Optimised and corrected parameter values based on soleus muscle data from Barclay et al. (1995). The EDL and soleus parameters were chosen to be equivalent. The corrected parameters were obtained through scaling the optimised parameters to account for typical recovery heat rates observed across a range of studies (Barclay, 2019) and to account for the temperature dependence of the maximum rate of oxidative phosphorylation (Davies & Tribe, 1969) to achieve reasonable energetic rates (Barclay & Weber, 2004).

| | $r_{rec} (J \mu\text{mol}^{-1})$ | $n_{ox}$ | $\phi_{ox,max} (\text{mmol s}^{-1})$ |
| --- | --- | --- | --- |
| Optimised parameters | 0.165 | 0.742 | 2.0255 |
| Corrected parameters | 0.0825 | 0.742 | 4.051 |

**Figure 2:**
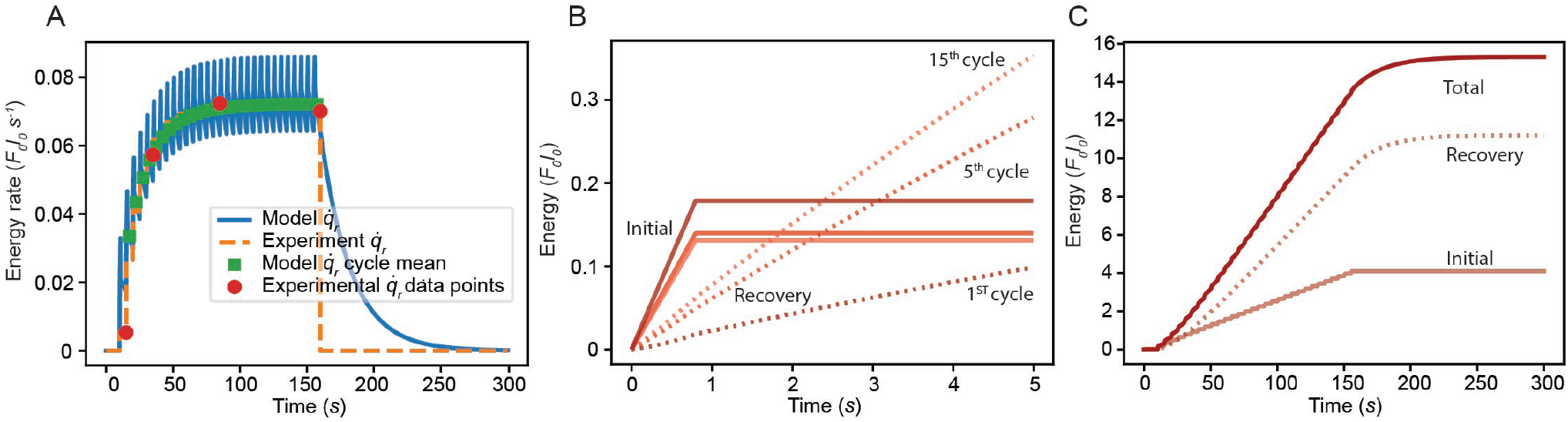
Optimisation of the bioenergetic parameters to the data from Barclay et al. (1995) for the soleus. The simulation protocol consisted of repeated contraction cycles of 5s with 800ms of tetanus contraction and 4.2s of rest. The initial energetic rates during the tetanus were computed from Barclay et al. (1995) Figure 3. A: Optimised model recovery heat rate 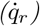 compared to the experimental data. Blue line is the time-varying recovery heat rate, orange dashed line is the interpolated recovery heat rate from the experimental data points (red dots). The model was fit to the experimental data by minimising the error between the interpolated experimental recovery heat rates and the cycle averaged model predicted energetic rates (green square). B: Energy use predicted by the model for the 1st (red), 5th (light red hue), and 15th (lightest red hue) contraction cycles. The initial energy (solid lines) given as input to the model and the corresponding predicted recovery heat (dotted line). C: The energy use predicted over the entire simulation of 30 contraction cycles and recovery period with total energy shown by the red solid line, the recovery heat shown by the transparent red dotted line, and the initial energy used as input given by the transparent solid line.

### 3.2 Validation

For an isometric contraction followed by a shortening phase (Fig. 3A,B), the model reproduced the time-course of energy consumption observed experimentally (Barclay & Weber, 2004) including both the temporal behaviour and magnitudes of the energetic rates (Fig. 3C,D, c.f. Barclay and Weber (2004) Figure 2B). The model achieved similar time constants of recovery, *≈* 11.5*s* for both soleus and EDL, to those observed experimentally Table 5, within 5% for soleus and 8% for EDL. The trend in model efficiencies showed higher initial efficiencies in the soleus compared to the EDL (*≈* 0.11), which follows the experimental trend of an *≈* 0.07 difference between the soleus and EDL (Table 5). Meanwhile, the total efficiencies (accounting for the initial energy and recovery costs) were within 0.05 between soleus in EDL, which was higher than the *≈* 0.009 observed experimentally (Table 5). The ratio of recovery to initial heat was also similar across muscles with *R* = 1.400 ± 0.003 for the soleus and *R* = 1.405 ± 0.008 for the EDL (Table 5)). The individual heat components demonstrated larger relative work and recovery heat contributions in the soleus compared to the EDL (Fig. 3E,F).

**Table 5:**
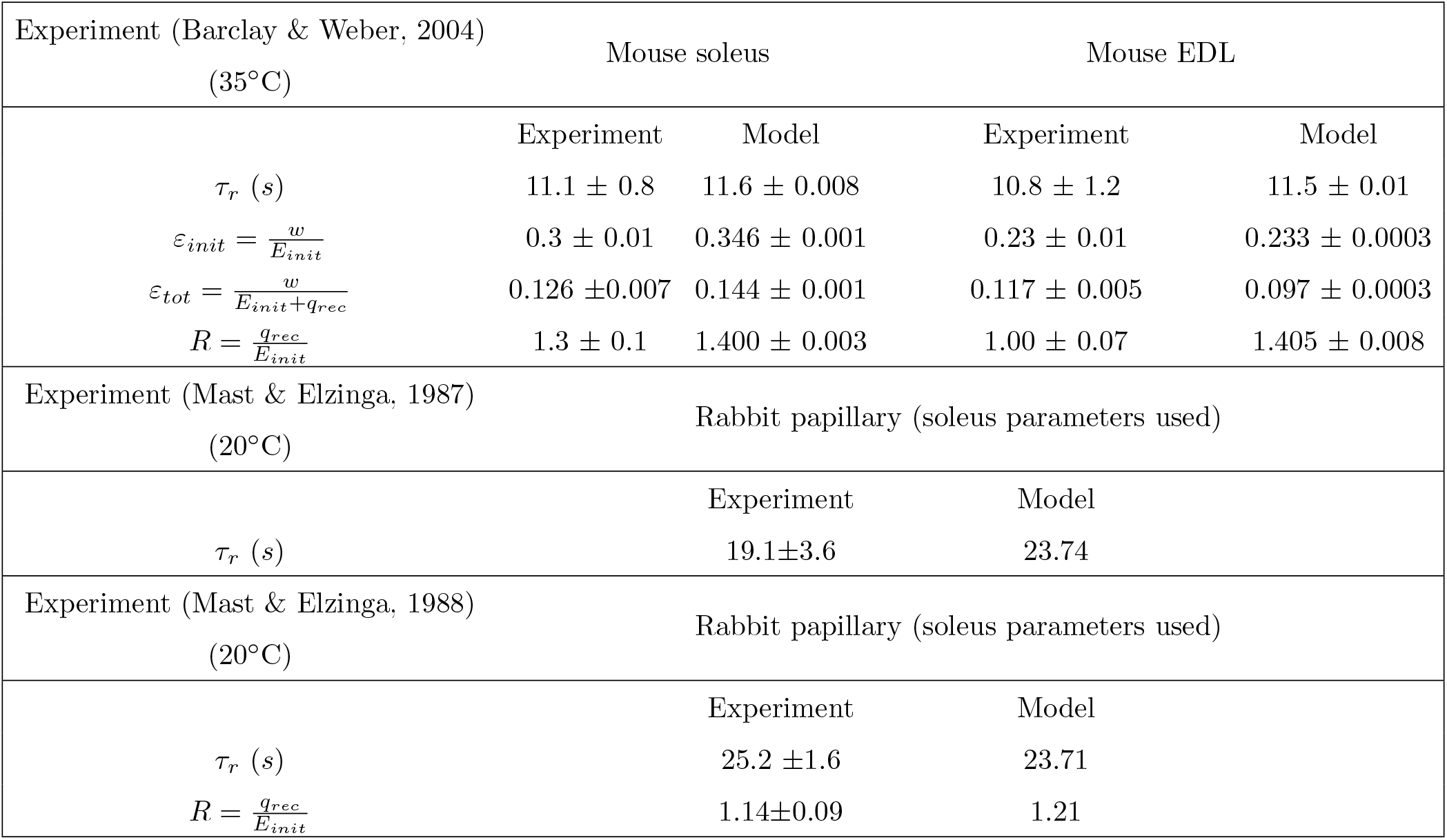
Comparison of time constants of recovery, efficiency values, and recovery-initial heat ratio between the model and experimental results from Barclay and Weber (2004), Mast and Elzinga (1987), and Mast and Elzinga (1988). τ_r_ is the time constant of the recovery processes obtained through fitting an exponential to the decay in recovery heat following muscle contractions. The efficiency values from Barclay and Weber (2004) are chosen for 35°C. ε_init_ represents the efficiency of the initial processes (not accounting for the recovery heat rates). ε_tot_ is the total efficiency accounting for both the initial and recovery processes. R is the ratio of recovery heat to the initial energy. Experimental and model values (when applicable) are Mean ± SEM. Slow-type muscle properties are used for the comparisons to rabbit papillary muscles used in Mast and Elzinga (1987) and Mast and Elzinga (1988). EDL: extensor digitorum longus.

**Figure 3:**
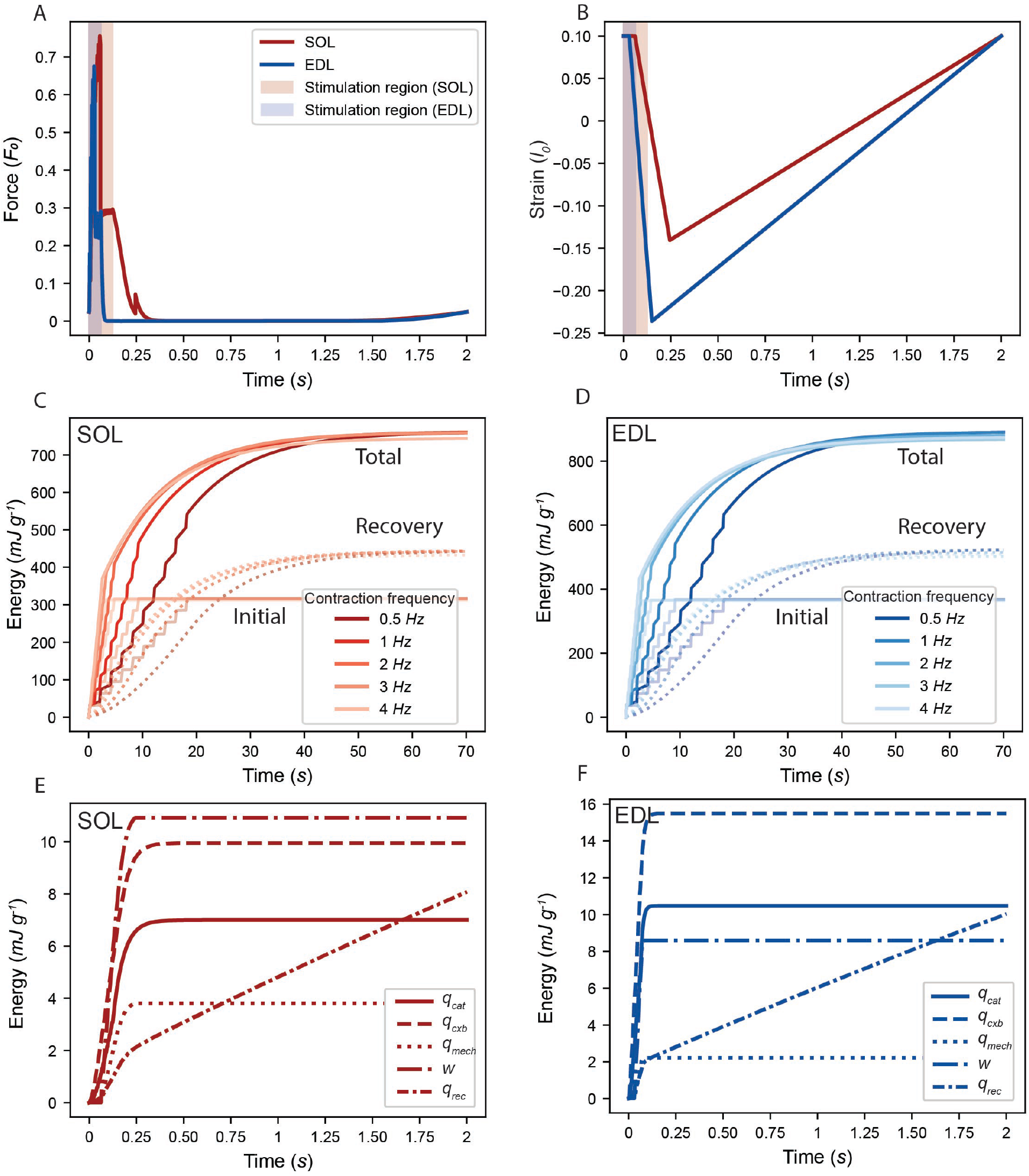
The mechano-bioenergetics model behaviour under experimental conditions from Barclay and Weber (2004). A,B: The experimental protocol for one contraction cycle including the stimulation range with force output (A) and muscle strain (B). C,D: Total energy consumed over the course of the simulation for various contraction frequencies for the soleus (C) and EDL (D). The total energy is given by the solid lines with each colour representing a different frequency. The transparent dotted lines give the recovery heat, while the transparent solid lines give the initial energy use. E,F: Relative components of the energetic rates for a single contraction cycle at 0.5Hz for the soleus (E) and EDL (F). q_cat_: Ca^2+^ transport heat, q_cxb_: cross-bridge heat, q_mech_: shortening/lengthening contribution, w: work, q_rec_: recovery heat, SOL: soleus, EDL: extensor digitorum longus.

The model parameter corrections (Table 4) were made based on time-constants observed in Barclay and Weber (2004), which no longer makes this an independent dataset to validate the time-constants obtained by the model. Thus, we compared with Mast and Elzinga (1987), which is independent and used a different measurement technique (*O*_2_ consumption) at 20^*°*^C (this allowed us to use the optimised time-constants directly as opposed to the corrected values Table 4). We observed a model time constant of 23.7*s*, which is within 20% of the experimental time constant of oxygen decay of 19.1±9.6*s* (Table 5). When we compared the normalised energetic rates at sub-maximal activation levels (stimulation frequencies between 40*Hz* and 160*Hz*), we observe similar behaviour to the experimental energetic rates with normalised (to the mean value) root mean square error *e*_*nrmse*_ = 10.20% for soleus and *e*_*nrmse*_ = 3.62% for EDL (Fig. 4). With consecutive twitches, we observe a decrease in Ca^2+^ transport heat that matches the trends from the experimental data set (Barclay, 2012) with *e*_*nrmse*_ = 18.66% and *e*_*nrmse*_ = 19.90% for soleus and EDL, respectively (Fig. 5). To ensure the proportion of recovery heat to initial heat was conserved at low stimulation frequencies, we compare to Mast and Elzinga (1988) and observe a ratio of initial to recovery heat of 1.21 which is within 6% of the experimental values of 1.14 ± 0.09 (Table 5). The time-course of energy consumption for the Mast and Elzinga (1987, 1988) are contained within Supplementary Material Figure S1.

**Figure 4:**
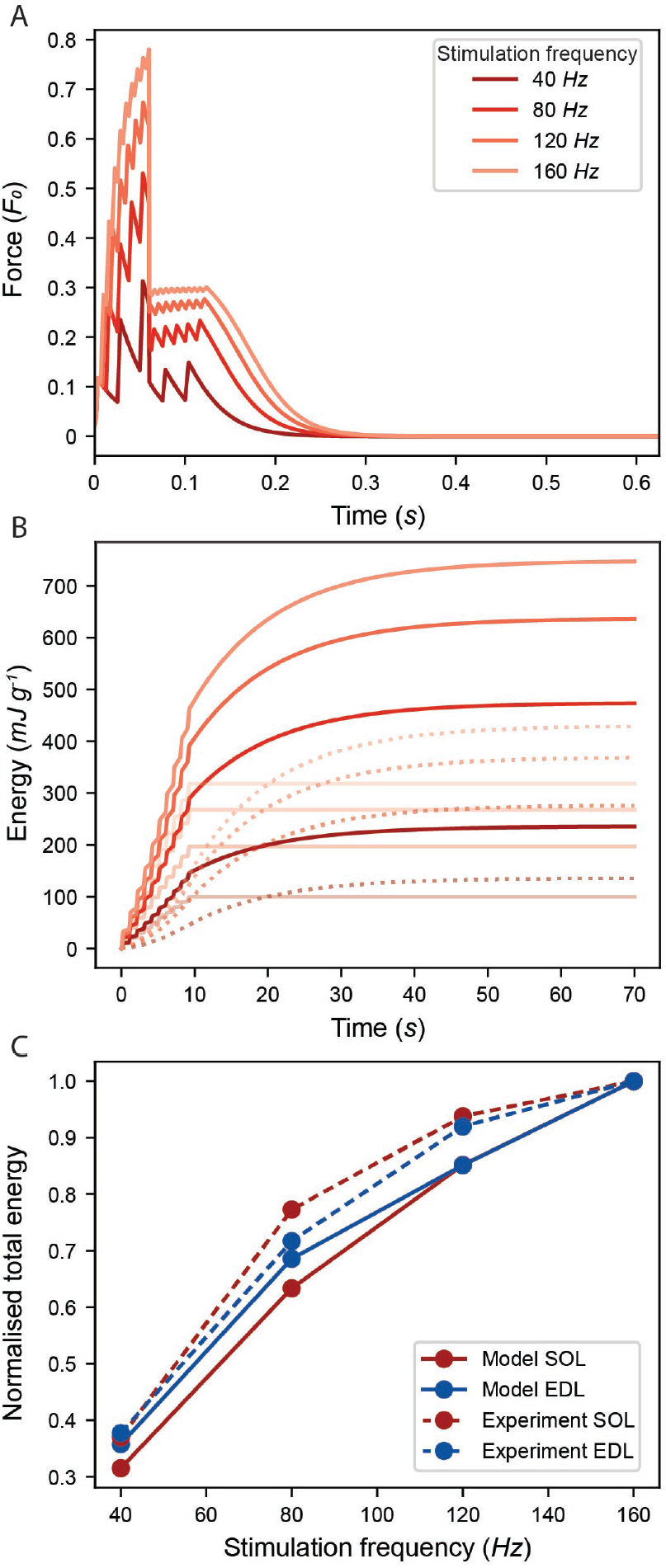
The mechano-bioenergetics model reproduces muscle energy rates across a ranges of stimulation frequencies. A: Example force produced over one contraction cycle with varying stimulation frequencies for the soleus. B: Example energy use over the whole simulation across different frequencies for the soleus. The solid lines correspond to total energy consumed, transparent solid lines represent initial energy use, and transparent dotted lines represent recovery energy. C: The normalised (to energy consumed at 160Hz) total energy consumed by the SOL and EDL across varying stimulation frequencies compared to experimental results from Lewis and Barclay (2014) (dashed lines). Note that errors in the magnitudes of the total energy consumed is expected given the difficulty in reproducing the precise experimental protocol; however the model is able to reproduce the dependence on stimulation frequency. SOL: soleus, EDL: extensor digitorum longus.

**Figure 5:**
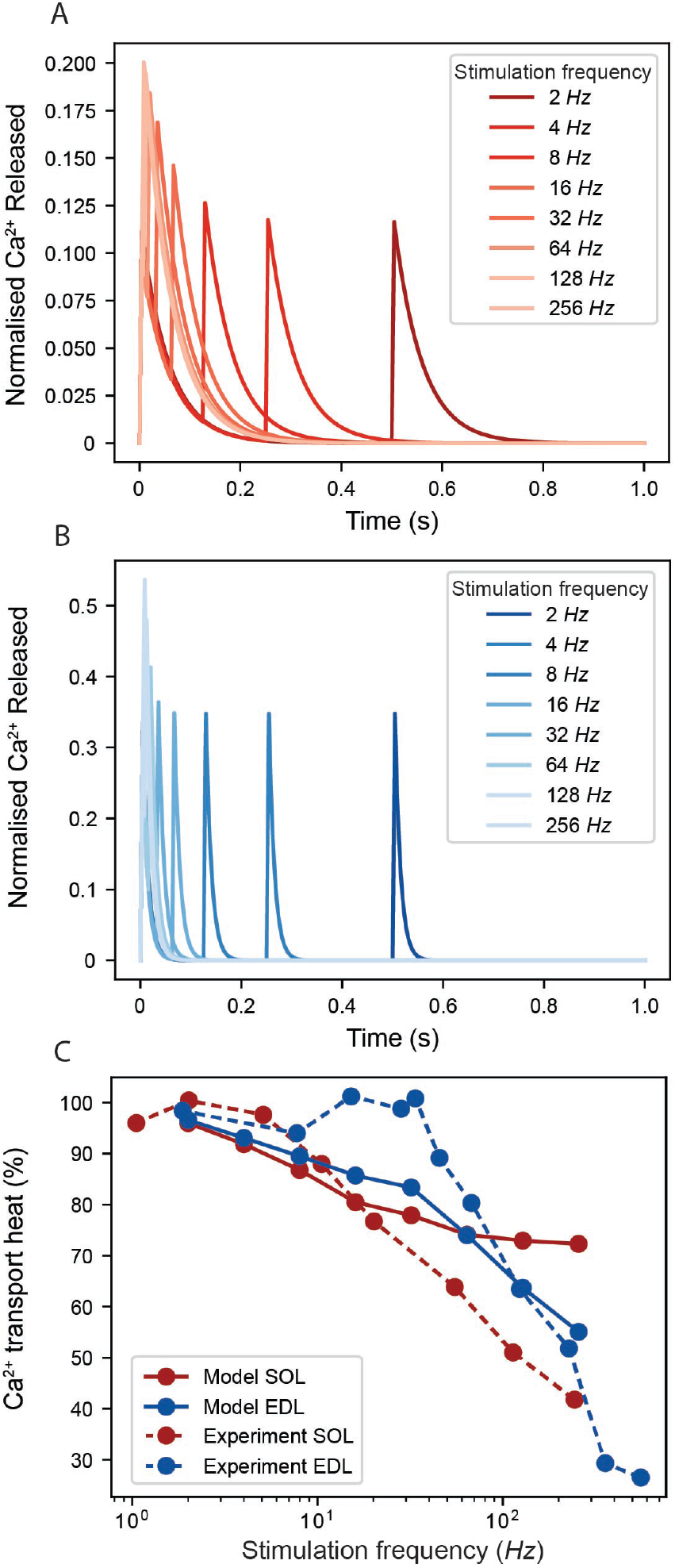
The mechano-bioenergetics model reproduces the dependence of Ca^2+^ transport heat on the frequency of stimulation. A,B: The Ca^2+^ released during two consecutive twitches at varying stimulation frequencies for the soleus (A) and the EDL (B). C: The Ca^2+^ transport heat of the second twitch as a percentage of the first twitch. Ca^2+^ transport heat rate followed the decrease with stimulation frequency observed in Barclay (2012) (dashed lines) in both the soleus and EDL. SOL: soleus, EDL: extensor digitorum longus.

## 4 Discussion

With parameters informed and validated on separate datasets, we found that the mechano-bioenergetics model predicted a similar time-course and magnitude of energetic rates to those observed experimentally (Barclay, 2012; Barclay & Weber, 2004; Lewis & Barclay, 2014; Mast & Elzinga, 1987, 1988). Further, it captured the time-dependence and recovery process not included in existing models. While the model did not necessarily capture the precise energy rates as in the validation studies, this was expected given the difficulty in reproducing experimental protocols as well as possible errors in the absolute heat measurements owing to challenges in calibration procedures (Woledge et al., 1985) and normalisation methods. We illustrated the performance of the mechano-bioenergetics model across a range of varied muscle fibre-types, contractile conditions, and stimulation intensities.

Energetic behaviour varies substantially across muscle fibre-types (Barclay, 1996), and capturing these differences is important, as fibre-types, which vary within muscles, individuals, species, and across the lifespan (Dick et al., 2026; Queeno et al., 2023), are closely related to muscle function (Goldspink, 1977). Across fibre-types, the model reproduced lower initial efficiencies in fast-type fibres compared to slow-type fibres (Barclay & Weber, 2004), which is expected given higher energetic demands of cross-bridge cycling in fast-type fibres (Barclay, 2019). While Barclay and Weber (2004) did not detect a difference in total efficiencies, we observed lower total efficiencies in the EDL - possibly a consequence of the same recovery parameters across fibres types. However, smaller changes in the total efficiency compared to initial efficiency across fibre-types were observed. The recovery time-constants were consistent across fibre-types, which is in agreement with Barclay and Weber (2004). The predicted recovery heat rates were comparable to experimental values, but likely overestimated as *R >* 1 in many simulations (Table 5); given the consistency of this overestimation, a scaling of *r*_*rec*_ to obtain an *R ≈* 1 could be made. The model demonstrated similar magnitudes of energetic rates between the fibre-types across contraction protocols (high shortening rate, low stimulation time for EDL and slow shortening rate, high stimulation time for soleus), which is consistent with experimental observations (Lewis & Barclay, 2014). Through capturing the the differences in fibre-type properties, we provide a model capable of making *in vivo* energy cost predictions, where fibre-type substantially influences energy use across a range of locomotor tasks (Swinnen et al., 2024).

Muscle contractions *in vivo* are often cyclical during locomotor tasks with contraction frequencies dependent on the speed of locomotion (Alexander, 1983). While obtaining muscle-level energetic rates *in vivo* is critical for accurately predicting energetics (Lichtwark et al., 2025), existing mathematical models (e.g., Lichtwark and Wilson (2005) and Umberger et al. (2003)) do not account for the time-dependent behaviour of muscle contraction and few models capture the recovery processes (Konno et al., 2025; Tsianos & MacFadden, 2016). These factors are particularly important for predicting the energetics of non-steady locomotion where the number, frequency, and intensity of the contractions vary. As observed in Fig. 3C,D, following a series of 10 contractions at slow frequencies, most of the recovery process takes place during the contractions, while at faster frequencies the recovery process occurs following the contraction. Thus, not accounting for recovery processes neglects these energetic rates completely, and without the time-dependent components of the energetics model its not possible to capture the dependence on contraction frequency.

This model was designed to be implemented at both the whole muscle level and, through validation of the model behaviour at low stimulation frequencies, the motor unit level. While we have informed the recovery model based on data from the mouse soleus (Barclay et al., 1995), the model would benefit from data on EDL recovery processes (to obtain fast-type muscle fibre properties) along with muscle energetic rates in the moderate frequency range of 5*Hz* to 40*Hz* (typical of *in vivo* motor unit firing rates (Enoka, 1995)). Here, we capture decreases in Ca^2+^ transport heat with decreases in simulation frequency, which is important for modelling the variable *in vivo* firing rates and is not captured with existing models. For example, with the increase in stimulation frequency from 2*Hz* to 256*Hz* in the soleus we saw a decrease in energy from 96% to 39% of the second twitch with this model (Fig. 5C), while the Konno et al. (2025) model predicted an increase in energetic rates (Supplementary Material Figure S2) - a consequence of only using information on the activation of the muscle and not the Ca^2+^ flow. While the model parameters presented here were informed by mouse data, the parameters in the bioenergetics model have been determined for human muscle (Vicini & Kushmerick, 2000) and could be used within this framework. Through accounting for physiological processes, the mechano-bioenergetics model is generalisable across muscles and contraction-types with the capability of implementation into whole-body musculoskeletal models and simulations of movement.

## 5 Author contributions

Conceptualisation: R.N.K., G.A.L., T.J.M.D.; Data curation: R.N.K.; Formal analysis: R.N.K.; Funding acquisition: G.A.L., T.J.M.D.; Investigation: R.N.K.; Methodology: R.N.K., G.A.L., T.J.M.D.; Software: R.N.K.; Supervision: G.A.L., T.J.M.D.; Validation: R.N.K.; Visualization: R.N.K.; Writing – original draft: R.N.K.; Writing review & editing: R.N.K., G.A.L., T.J.M.D.

## 6 Acknowledgments

This work is supported by an Australian Research Council Discovery Project Grant (DP230101886) to T.J.M.D. G.A.L. is supported by an Australian Research Council Future Fellowship (FT190100129). R.N.K. is supported by a Natural Science and Engineering Research Council of Canada Postgraduate Scholarship and by the Commonwealth through an Australian Government Research Training Program Scholarship DOI: https://doi.org/10.82133/C42F-K220.

