## Supplementary Material for "A time-dependent mechano-bioenergetics model of muscle contraction"

### Supplementary Material: A time-dependent mechano-bioenergetics model of muscle contraction

Konno, R. N.[1,*], Lichtwark, G. A.[2,3], and Dick, T. J. M.[1]

[1]*School of Biomedical Sciences, The University of Queensland, Australia*
[2]*School of Exercise & Nutrition Sciences, Queensland University of Technology, Australia*
[3]*School of Human Movement & Nutrition Sciences, The University of Queensland, Australia*

#### S1 Hill-type model formulation

We assume that the fraction of $Ca^{2+}$ bound to troponin is proportional to the muscle activation level such that

$$\hat{a} = \frac{c_{catn}}{c_{CaTn,max}}, \tag{S1}$$

where $c_{CaTn,max}$ is the maximum amount of bound Troponin. The normalised force produced by the muscle can then be calculated based on a Hill-type model

$$\hat{F}_m = \hat{a}\,\hat{F}_{ce,l}(\epsilon_{ce})\,\hat{F}_{ce,v}(\dot{\epsilon}_{ce}) + \hat{F}_{pe}(\epsilon_{ce}). \tag{S2}$$

$\epsilon_{ce}$ is the strain in the muscle and $\dot{\epsilon}_{ce}$ is the strain rate. $\hat{F}_{ce,l}(\epsilon)$ gives the force-strain relationship (Gordon et al., 1966)

$$\hat{F}_{ce,l}(\epsilon_{ce}) = e^{-|\frac{\epsilon_{ce}^{s_k}-1}{w}|^r}. \tag{S3}$$

$s_k$, $w$, and $r$ are constants governing the shape of the force-strain relationship. $\hat{F}_{ce,v}(\dot{\epsilon}_{ce})$ gives the force-strain-rate relationship (Hill, 1938)

$$\hat{F}_{sr}(\dot{\epsilon}) = \begin{cases} \frac{1+\left(\frac{\dot{\epsilon}_{ce}}{\dot{\epsilon}_{ce,max}}\right)}{1-\left(\frac{\dot{\epsilon}_{ce}}{\kappa\dot{\epsilon}_{ce,max}}\right)}, \text{if} \quad \dot{\epsilon}_{ce} \leq 0, \\ 1.5 - 0.5\,\frac{1-\left(\frac{\dot{\epsilon}_{ce}}{\dot{\epsilon}_{ce,max}}\right)}{1+\left(\frac{7.56\,\dot{\epsilon}_{ce}}{\kappa\dot{\epsilon}_{ce,max}}\right)}, \text{if} \quad \dot{\epsilon}_{ce} > 0, \end{cases} \tag{S4}$$

where $\dot{\epsilon}_{ce,max}$ is the maximum shortening rate of the muscle and $\kappa$ is a parameter governing the curvature of the force-strain-rate relationship. $\hat{F}_{pe}(\epsilon_{ce})$ gives the passive force length relationship accounting for any parallel passive structures within the muscle (connective tissue, titin, etc.)

$$\hat{F}_{pe}(\epsilon_{ce}) = \begin{cases} 2.64\epsilon_{ce}^2 - 5.3\,\epsilon_{ce} + 2.66, \text{ if } \epsilon_{ce} > 0, \\ 0, \text{ otherwise.} \end{cases} \tag{S5}$$

#### S2 Optimisation functions

Optimisation functions used to determine the initial energetic cost. $\mathcal{L}_{cxb}$ and $\mathcal{L}_{cat}$ enforce the cross-bridge and $Ca^{2+}$ transport heat components, respectively. $\mathcal{L}_{short}$ enforces the mechanical dependence on the shortening rate. $\mathcal{L}_{ratio}$ enforces ratio in $Ca^{2+}$ transport heat to total initial heat across stimulation frequencies.

$$\mathcal{L}_{cxb} = (0.6\,\dot{q}_{exp,iso} - \dot{q}_{cxb,iso})^2, \tag{S6a}$$

$$\mathcal{L}_{cat} = (0.4\,\dot{q}_{exp,iso} - \dot{q}_{cat,iso})^2, \tag{S6b}$$

$$\mathcal{L}_{short} = \frac{1}{|\dot{q}_{exp}|}||\dot{\mathbf{q}}_{exp} - \dot{q}_{exp,iso} - \dot{\mathbf{q}}_{short}||^2, \tag{S6c}$$

$$\mathcal{L}_{ratio} = ||\mathbf{q_{cat,iso}} - 0.4\,(\mathbf{q_{cat,iso}} + \mathbf{q_{cxb,iso}})||^2\,. \tag{S6d}$$

$\dot{q}_{exp,iso}$ represents the experimental isometric heat rate. $\dot{q}_{cxb,iso}$ and $\dot{q}_{cat,iso}$ represent the mean cross-bridge and $Ca^{2+}$ transport heat rates evaluated at zero shortening rate. $\dot{\mathbf{q}}_{exp}$ is the vector of experimentally measured experimental heat rates and $\dot{\mathbf{q}}_{short}$ is the vector of model shortening heat rates, with entries corresponding to a given shortening rate. $|\dot{q}_{exp}|$ is the size of $\dot{q}_{exp}$ and $||x||$ represents the 2-norm of a given vector $x$. The heat rates $\dot{q}_{cxb,iso}$, $\dot{q}_{cat,iso}$, and $\dot{\mathbf{q}}_{short}$ were averaged over the shortening period of the contraction. $\mathbf{q_{cat,iso}}$ and $\mathbf{q_{cxb,iso}}$ gives the vector of $q_{cat,iso}$ and $q_{cxb,iso}$ computed over stimulation intensities of 40, 60, 120, and $160Hz$.

### S3 Simulation protocols

This section contains the detailed simulation protocols for the numerical simulations performed within this study. The implementations of these specific protocols are available at https://github.com/ryankonno/TimeDependentEnergetics.

#### S3.1 Barclay et al., 2010 protocol

The optimisation simulations follow the protocol from (Barclay et al., 2010) with stimulation frequencies of $150Hz$. The protocol consists of an isometric phase followed by muscle shortening at a fixed rate. The shortening rate was between 0 and 5 $l_0\, s^{-1}$. The muscle was assumed to be shortening over the plateau of the force-length relationship, so these effects were ignored. Muscle energetic rates were measured as the mean rates over the shortening period of the contraction.

#### S3.2 Barclay et al., 1995 protocol

This protocol consisted of cyclical isometric contractions with a contraction and rest phase. The rest phase gives solely recovery heat, while the contraction phase gives a combination of the recovery and initial heat. We prescribe the initial heat rate as input to the model. This was obtained from the digitised absolute heat measures as the slope of the heat consumed during the contraction, subtracted from the heat consumed during the rest period (Lichtwark et al., 2025). The simulation was performed for 150 $s$ (30 cycles at 5 $s$ per cycle) with the muscle stimulated for 0.8 $s$ per cycle.

#### S3.3 Barclay and Weber, 2004 protocol

The simulation protocols are designed to follow the experimental protocols, however, the experimental procotocols were designed to optimise power output for the specific muscle preparations, and the specific parameters are not available within the text. The muscle contraction cycles were performed at contraction frequencies of 0.5 to 4 $Hz$ with muscle stimulated for a fixed time period held constant across frequencies ($0.125s$ for soleus and $0.063s$ for EDL). Each contraction cycle consisted of an isometric phase at the start of stimulation ($0.06s$ for soleus and $0.03s$ for EDL), then an isokinetic shortening phase at 1.3 $l_0\, s^{-1}$ for the soleus and 2.8 $l_0\, s^{-1}$ for the EDL (based on values in (Barclay & Weber, 2004)). Shortening continued until muscle was deactivated, then the muscle was lengthened back to its original length. The original starting length was chosen to be $0.1\, l_0$ to allow muscle to shorten over the plateau of the force-length relationship. The total cycle length is determined based on the cycle frequency with 10 cycles at each frequency. The simulation was performed for $70s$ to allow for muscle recovery. The muscles were stimulated at frequencies of $150Hz$ and $175Hz$ for the soleus and EDL, respectively.

#### S3.4 Lewis and Barclay, 2014 protocol

These simulations followed the same protocol as in (Barclay & Weber, 2004), but were only performed at $1Hz$ frequency. For this protocol the stimulation frequencies were varied from $40Hz$ to $160Hz$.

#### S3.5 Mast and Elzinga, 1987 protocol

This protocol consisted of a series of 120 muscle twitches at a frequency of $0.5Hz$. The simulation was performed for a total length of $300s$. Muscle was held isometric for the duration of the experiment. Soleus muscle properties were chosen for these experiments with the optimised parameters for 20°C to match experimental temperatures.

#### S3.6 Mast and Elzinga, 1988 protocol

This protocol consisted of a series or 10 twitches at $0.2Hz$ frequency. The simulation was performed for a total length of $100s$. Muscle was held isometric for the duration of the experiment. Soleus muscle properties were chosen for these experiments with the optimised parameters for 20°C to match experimental temperatures.

#### S4 Time-varying energetic rates

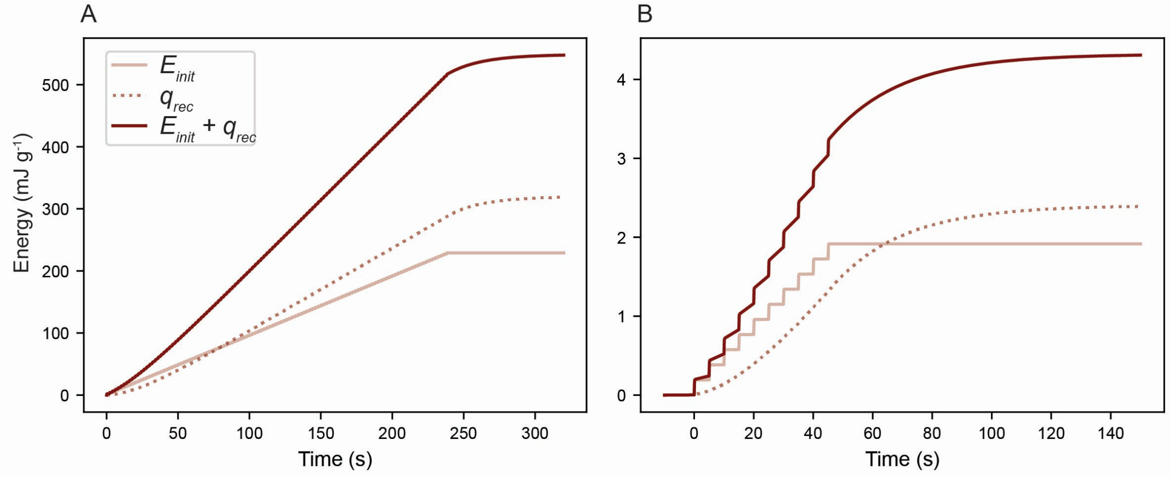


***Figure S1: Time-course of energy use during the Mast and Elzinga (1987) (A) and Mast and Elzinga (1988) (B) simulations.*** *Soleus parameters are used for the simulations. Solid line represents total energy use, transparent solid line represents initial energy use, and transparent dotted line represents recovery heat.*

#### S5 Cross-model comparison

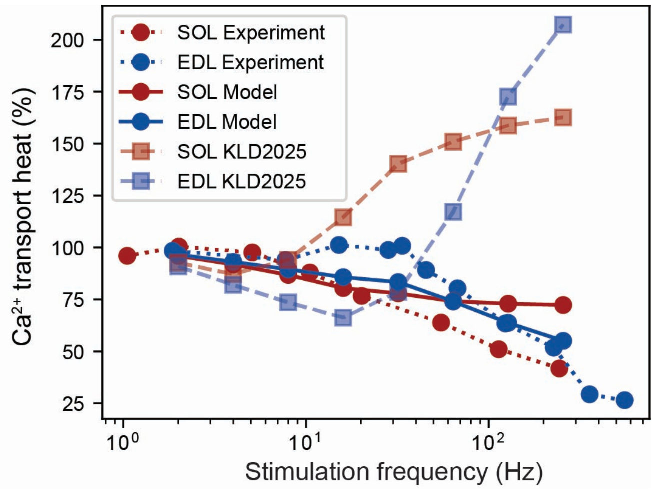


***Figure S2: Comparison of model results in Figure 5C to the KLD2025 model (Konno et al., 2025).*** *We observe an an increase in energetic rates in the model developed by Konno et al. (2025) at high frequencies in contrast to the decrease observed in the current model. The $Ca^{2+}$ transport heat for the KLD2025 model was computed from the combined activation and maintenance heat by assuming 40% was contributed to the $Ca^{2+}$ transport costs. SOL: soleus; EDL: extensor digitorum longus.*
